# UV-B exposure and exogenous hydrogen peroxide application leads to cross-tolerance toward drought in *Nicotiana tabacum* L

**DOI:** 10.1101/2021.02.26.432958

**Authors:** O Diana Sáenz-de la, Luis O. Morales, Åke Strid, Irineo Torres-Pacheco, Ramón G. Guevara-González

## Abstract

Acclimation of plants to water deficit involves biochemical and physiological adjustments. Here, we studied how UV-B exposure and exogenously applied hydrogen peroxide (H_2_O_2_) potentiates drought tolerance in tobacco (*Nicotiana tabacum* L.). Separate and combined applications for 14 days of 1.75 kJ m^−2^ day^−1^ UV-B radiation and 0.2 mM H_2_O_2_ were assessed. Both factors, individually and combined, resulted in inhibition of growth. Furthermore, the combined treatment led to the most compacted plants. UV-B- and UV-B+H_2_O_2_-treated plants increased total antioxidant capacity and foliar epidermal flavonol content. H_2_O_2_- and UV-B+H_2_O_2_-pre-treated plants showed cross-tolerance to a subsequent 7-day drought treatment. Plant responses to the pre-treatment were notably different: i) H_2_O_2_ increased the activity of catalase, phenylalanine ammonia lyase and peroxidase activities, and ii) the combined treatment induced epidermal flavonols which were key to drought tolerance. We report synergistic effects of UV-B and H_2_O_2_ on transcription accumulation of *UV RESISTANCE LOCUS 8, NAC DOMAIN PROTEIN 13* (*NAC13*), and *BRI1-EMS-SUPPRESSOR 1* (*BES1*). Our data demonstrate a pre-treatment-dependent response to drought for *NAC13, BES1* and *CHALCONE SYNTHASE* transcript accumulation. This study highlights the potential of combining UV-B and H_2_O_2_ to improve drought tolerance which could become a useful tool to reduce water use.

## Introduction

Extreme weather events limit plant production, which, in turn, may affect agricultural important plant species so as to reduce food production. In addition, the frequency of such extreme events are likely to increase as climate change worsens (Lesk, Rowhani & Ramankutty 2016). Therefore, plant drought tolerance is an important trait which we need to more completely understand at the physiological and molecular level (Godfray et al., 2010). Drought stress diminishes crop growth, disturbs plant water and plant nutrient relations, reduces photosynthesis and causes oxidative damage due to the generation of reactive oxygen species (ROS) (Salehi-Lisar & Bakhshayeshan-Agdam, 2016). Thus, plant drought tolerance is a complex process that mainly involves osmotic adjustment, osmoprotection, and antioxidative activity in the form of a ROS scavenging defense system (Farooq, Wahid, Kobayashi, Fujita & Basra, 2009). In addition, some chemical and physical stressors, that at high concentrations cause toxicity, may at low or moderate concentrations play a positive role by pre-conditioning plants against the negative influence of other abiotic factors, such as drought (Vázquez-Hernández et al., 2019). Such pre-conditioning is often refered to as cross-tolerance.

The ultraviolet region of the electromagnetic spectrum is conventionally defined as UV-C (<280 nm), UV-B (280-315 nm) and UV-A (315-400 nm) (Björn, 2015). Approximately 95% of UV radiation reaching the Earth is UV-A, whereas UV-C and UV-B radiation below 290 nm are absorbed by ozone in the stratosphere (NASA, 2018). UV-B, despite of being only a small proportion of the UV spectrum, has the potential of altering plant morphology and metabolism (Jenkins, 2009). Previous studies have reported that UV-B stress has negative effects on photosynthesis, plant growth and biomass accumulation (Jansen, Gaba & Greenberg 1998; Jordan, 2002; Frohnmeyer & Staiger, 2003; Albert, Mikkelsen, Ro-Poulsen, Arndal & Michelsen 2011). However, research of the last decades has shown that ambient levels of UV-B, perceived through the UV-B photoreceptor UV RESISTANCE LOCUS 8 (UVR8), regulate photomorphogenesis and defense responses, including photorepair processes, antioxidant activities and UV-screening (Jenkins, 2009; Rizzini et al., 2011; Jansen & Bornman, 2012; Morales et al., 2013). In this sense, UV-B radiation has been suggested to act as a signal that can promote tolerance to biotic and abiotic stress conditions. The potential use of UV-B radiation to alleviate the effects of drought stress has been reported in different plant species such as silver birch (Robson, Hartikainen & Aphalo, 2015a), rice (Dhanya Thomas, Dinakar & Puthur, 2020) and wheat (Kovács et al., 2014). Plant responses to the combination of drought and UV-B depend on whether the treatments are simultaneous or sequential (Escobar-Bravo et al., 2019). The sensitivity of the plant species also determines whether the interaction between drought and UV-B is additive, synergistic or antagonistic (Bandurska & Cieślak, 2013). Cross-tolerance to drought may be activated by defense-related molecules, such as hydrogen peroxide (H_2_O_2_), nitric oxide (NO), abscisic acid (ABA), ethylene, jasmonic acid (JA) or salicylic acid (SA), which are produced as responses to UV-B (A.-H.-Mackerness, John, Jordan & Thomas, 2001; Bandurska & Cieślak, 2013; Tossi, Lamattina, Jenkins & Cassia, 2014; Mannucci et al., 2020). However, how UV-B mechanistically interact with these molecules while inducing cross-tolerance to drought remains poorly understood.

Plant responses to H_2_O_2_ have received much attention because of the role of H_2_O_2_ as a signaling molecule, being considered an elicitor that, when exogenously applied, can induce defense in plants prior to stress exposure (Vazquez-Hernandez et al., 2019; Parola-Contreras et al., 2020). Among H_2_O_2_-triggered responses, cross-tolerance to abiotic and biotic stresses is often observed (Hossain et al., 2015). Alleviation of drought using H_2_O_2_-controlled elicitation has been reported in cucumber (Sun, Wang, Liu & Peng, 2016), rice (Sohag et al., 2020) and soybean (Ishibashi et al,. 2011). Drought stress increases levels of ROS, such as H_2_O_2_ and singlet oxygen (O_2_), in chloroplast, peroxisome, and mitochondria, which in turn can lead to oxidative damage (Cruz De Carvalho, 2008). Despite this, priming with H_2_O_2_ can protect plants against damage by triggering antioxidant enzymes, such as superoxide dismutase (SOD), catalase (CAT), glutathione peroxidase (GPX) and ascorbic acid peroxidase (APX), thus reducing accumulation of different ROS (Hossain et al., 2015). This means that oxidative damage may be effectively alleviated by the antioxidant machinery after activation through exogenous H_2_O_2_ application. Application of either UV-B in barley or H_2_O_2_ in cucumber provided drought protection (Bandurska, Niedziela & Chadzinikolau, 2013; Sun et al., 2016). However, in what way UV-B and H_2_O_2_ together can prime drought tolerance remains unknown.

The aim of this study was to evaluate cross-tolerance responses under moderate drought conditions in *Nicotiana tabacum* plants. A factorial experiment was implemented to assess the interaction of simultaneous exposure to UV-B exposure and exogenous application of H_2_O_2._

## Materials and methods

### Plant material, growth conditions

Tobacco seeds (*Nicotiana tabacum* L. cv. xanthi nc) were surface sterilized, sown *in vitro* on Murashige and Skoog medium and kept in a germination chamber at 25 ± 1°C, 60 ± 5% of relative humidity (RH) and 70-85 µmol m^-2^ s^-1^ of photosynthetically active radiation (PAR). After 10 days from sowing, seedlings were transplanted into individual plastic pots containing fertilized peat moss-based substrate. Additionally, 25% diluted Hoagland solution was applied one week after transplanting and during the rest of the experiment to compensate for the loss of nutrients in the substrate (See Supplementary Table S1). Irrigation was carried out by adding water to the trays containing the pots. Plants were then grown in a greenhouse at 25°C day/ 20°C night and 80% RH. Seedlings were exposed to PAR from high-pressure sodium lamps (Vialox NAV-T Super 4Y; Osram, Johanneshov, Sweden) corresponding to 120–240 μmol m^−2^ s^−1^ for 16 h per day (from 06:00 h to 22:00 h) daily.

### Experimental treatment conditions

When plants were three weeks old, UV-B- and H_2_O_2_-pre-treatments were applied for 14 days, resulting in four plant treatment groups: control, UV-B-, H_2_O_2_- and UV-B+H_2_O_2_-pre-treated plants. The pre-treatment conditions were as follows:

i. Supplementary UV-B was applied for 4 h per day (from 10:00 h to 14:00 h). UV-B irradiation was provided by fluorescent lamps (Philips TL40/12 UV, Eindhoven, The Netherlands) filtered through 0.13 mm cellulose acetate sheets (Nordbergs Tekniska AB, Vallentuna, Sweden) to remove any UV-C radiation, and normalized to 300 nm (Yu & Björn 1997; Kalbina, Li, Kalbin, Björn & Strid, 2008). 122 mW m^−2^ of plant-weighted UV-B was applied for 4 h giving a total irradiation of 1.75 kJ m^−2^ day^−1^. UV radiation was measured using an Optronic Laboratories OL756 (Orlando, FL) double monochromator spectroradiometer. Control plants were exposed to white light only in the same chamber as the UV-B plants by blocking UV radiation using Perspex (Plastbearbetning AB, Norsborg, Sweden).
ii. H_2_O_2_ (0.2 mM) was foliar sprayed to dew point (using 2.5 mL per plant) and applied to irrigation (using 10 mL per plant). H_2_O_2_ was applied every third day giving a total of 5 applications.
iii. Plants were exposed to a combined treatment of UV-B and H_2_O_2_, under the aforementioned conditions for each factor.

Drought treatment started at day 15, when plants were five weeks old. To achieve moderate drought conditions, pots were weighed at 100% soil water content and set to 60% of field capacity by estimating a 40% reduction in pot weight. After applying water withdrawal for 3 days, soil moisture reached 40-45% of field capacity. Drought treatment was accomplished by watering again the plants so that field capacity remained at 40-45%. Viability was determined by visual assessment and measuring leaf relative water content (RWC). The RWC (%) was measured in the second fully expanded leaf from the apex. Leaves were collected, the fresh weight (Fw) was recorded, and the leaves incubated in distilled water for 4h at 4°C in the dark. The leaves were blotted, and the turgid weight (Tw) measured. Finally, leaves were dried at 80°C overnight and the dry weight (Dw) measured. RWC was calculated as RWC= (Fw - Dw) / (Tw - Dw) (Jones 2007).

For further analysis, leaf samples were collected from the top parts of the plants during the 21 days of the experiment at 0, 4, 28 h and at days 7, 14 and 21. Root samples were collected at the end of the drought treatment (day 21). Samples were frozen in liquid nitrogen, stored at −80°C, and freeze-dried for 3 days (Lyolab 3000, ThermoFisher Scientific). Three replicates per sampling point were collected for all the analyses.

### Morphological measurements and non-invasive leaf epidermal flavonol content

Morphological parameters were measured in six plants per treatment at the beginning (day 1), middle (day 7), and end of the pre-treatment (day 14). Stem length was measured using a ruler, the number of leaves was counted, and the stem diameter was measured using a digital caliper. The adaxial side epidermal flavonol index of young leaves was measured at hour 0, 4, and 28, and every third day thereafter until the end of the experiment using a DUALEX® SCIENTIFIC optical sensor (ForceA, France).

### Total antioxidant capacity

Total antioxidant capacity was measured using a commercially available kit (Total Antioxidant Capacity Assay kit, Sigma, St Louis). 100 mg of leaf tissue from a pool of leaves was frozen at −80°C and homogenized using liquid nitrogen and a mortar and pestle. Samples were extracted in 1 mL of 4°C 1 X Phosphate Buffered Saline and the supernatant was diluted 1:30 to bring values within the range of the kit standards. Samples were assayed according to the manufacturer’s protocol, by comparing the absorbances of diluted extracts at 570 nm with Trolox standards. Values were then normalized to tissue fresh weight as equivalents of mg Trolox using the standard reference curve.

### Preparation of leaf extracts and enzymatic assays

Lyophilized leaf samples (0.05 g) were ground with 2 mL ice cold potassium-phosphate buffer (0.05 M, pH 7.8) using a mortar and pestle. Samples were centrifuged at 12,000 rpm for 15 min at 4°C. The supernatants were stored at −20°C until further used for assays of CAT, PAL, SOD and POD activities. Total soluble protein content was determined spectrophotometrically (λ595nm) according to the standard Bradford assay (Bradford, 1976), using bovine serum albumin as standard.

The catalytic activity of catalases (CAT; EC 1.11.1.6) was measured spectrophotometrically (λ240nm) by monitoring the rate of H_2_O_2_ decrease for 6 minutes at room temperature as described by Afiyanti & Chen (2014). The reactions were started by adding 1.4 mL 50 mM potassium phosphate buffer (pH 8.0), 140 µL 100 mM H_2_O_2_, and 70 µL sample extract. Catalase activity was expressed in units of mmol of H_2_O_2_ decomposed (mg protein * min)^-1^.

Phenylalanine ammonia-lyase (PAL; EC 4.3.1.5) activity was determined by measuring spectrophotometrically (λ290nm) the increases in the formation of cinnamic acid according to Toscano, Ferrante, Leonardi & Romano (2018) by incubating 0.1 mL of sample extract in 1.5 mL of 0.1 M borate buffer (pH 8.8) containing 10 mM L-phenylalanine for 1 h at 40°C. The reaction was stopped by the addition of 0.25 mL 1 N HCl. One unit (U) of PAL released 1 μmol of cinnamic acid (min)^-1^ at pH 8.8 and 40°C. The PAL activity was expressed as U (mg protein)^-1^.

The superoxide dismutase activity (SOD; EC 1.15.1.1) was assayed spectrophotometrically (λ560nm) using the method of Hayat, Ahmad, Ali, Ren & Cheng (2018). 50 μL of the sample extract was added to 2.95 mL of the reaction mixture. The mixture contained 1.5 mL 0.05 M phosphate buffer (pH 7.8), 0.3 mL 0.1 mM EDTA, 0.3 mL 0.13 M methionine, 0.3 mL 0.75 mM nitroblue tetrazolium (NBT), 0.3 mL 0.02 mM riboflavin and 0.25 mL distilled water. One unit (U) of SOD activity was defined as the amount of enzyme required for 50% inhibition of photochemical reduction of NBT, expressed as U (mg protein)^-1^.

Peroxidase activity was recorded using a commercially available kit (Amplex® Red Hydrogen Peroxide / Peroxidase Assay Kit, ThermoFisher Scientific). Samples were diluted 1:5 to bring values within the range of the kit standards. Samples were assayed according to the manufacturer’s protocol, by detecting the fluorescence signal in a plate reader with emission at 590 nm.

### Determination of proline content in the roots

Proline was measured in roots by the ninhydrin-based colorimetric method as described by Lee, Kim, Park, Choi & Lee (2018). Lyophilized root tissue (0.05 g) was extracted by homogenization in 1 mL of 1% (*w*/*v*) sulfosalicylic acid. One mL of plant extract was reacted with 2 mL acidic ninhydrin (1.25% [*w*/*v*] ninhydrin in 80% [*v*/*v*] acetic acid) and incubated at 100°C for 60 min. The reaction was stopped by putting the samples in an ice bath for 10 min. The absorbance was measured using a spectrophotometer at λ510 nm, and the proline concentration was calculated using the standard curve.

### Gene expression analysis

RNA extraction was performed using Direct-zol RNA miniprep kit (BioAdvanced Systems Co.) according to the manufacturer’s instructions. cDNA synthesis was performed using the Maxima First Strand cDNA Synthesis (ThermoFisher Scientific) for RT-qPCR according to the instructions of the provider (10 min at 25°C followed by 15 min at 50°C). Accumulation of mRNA transcripts for marker genes involved in UV-B signaling or responding to UV-B (*UVR8*; *BRI1-EMS-SUPPRESSOR 1, BES1*; *CHALCONE SYNTHASE, CHS*; *NAC DOMAIN PROTEIN 13, NAC13*) was measured. The following forward (F) and reverse (R) primers, designed by using the Primer-Blast tool from the National Center for Biotechnology Information, were used in the experiments: UVR8F - GCGGACCATGCAGAGATGC; UVR8R – CACAAGCCTAGGTGATGTCTGT; NAC13F – GCGCCTGGTACAGGTGATT; NAC13R - GCATATGCTGGCTTTGCAGG; BES1F - ATCGCAAGGGACACAAGCC; BES1R - CCAACTCGAGAAGGACTCGG; CHSF – TTCTCCGATTGGCCAAGGAC; CHSR - CACTTGGGCCACGAAATGTG.

Quantitative PCR was performed using the Maxima SYBR Green/ROX qPCR Master Mix (Thermo Scientific) kit following the manufacturer’s instructions. The PCR conditions were as follows: 5 min at 94°C, 40 cycles of 1 min at 94°C, and 1 min at 56°C for *UVR8*, 64°C for *NAC13*, 62°C for *UVR8* and *CHS*. Bio-Rad CFX manager software was used to automatically calculate the cycle threshold (Ct) value for each reaction. Each reaction was performed at least in duplicate in two independent experiments. The expression of all genes was normalized to the mean of the housekeeping gene *elongation factor 1 alpha* (*EF-1A*). The primers for EF-1A were: forward – TGAGATGCACCACGAAGCTC; reverse – CCAACATTGTCACCAGGAAGTG. The quantification of mRNA levels was based on the relative quantification method (2^−ΔΔCt^) (Livak & Schmittgen, 2001).

### Statistical analysis

Statistical analyses were carried out using the GraphPad prism 9.0 program (GraphPad Software, San Diego, CA). Data were analyzed by using two-way ANOVA followed by Tukey’s test at p≤0.05 except for gene expression data where one-way ANOVA was used. Data are presented as the mean ± standard deviation of six replicates for morphological parameters and epidermal flavonol content estimated with Dualex, and of three replicates for TEAC, enzymatic activities, RWC and proline content. Gene expression data are presented as the mean ± standard error of at least two replicates.

## Results

### Effects of pre-treatments on morphological and biochemical features

Among the morphological parameters studied, UV-B and H_2_O_2_ treatments individually reduced stem length, resulting in plants shorter than those in the control group (Fig. 1A). The UV-B+H_2_O_2_ treatment led to a reduced stem length compared to control and to the individual effect of the H_2_O_2_ treatment (Fig. 1A). The basal stem diameter was only decreased by the combined UV-B+H_2_O_2_ treatment (Fig. 1B). In addition, the number of leaves of UV-B+H_2_O_2_-treated plants was lower compared to the control and to the effect of either of the individual factors (Fig. 1C). Thus, the combined treatment effect on plant morphology was shorter and more compact plants (Fig. 2A).

**Figure 1.**
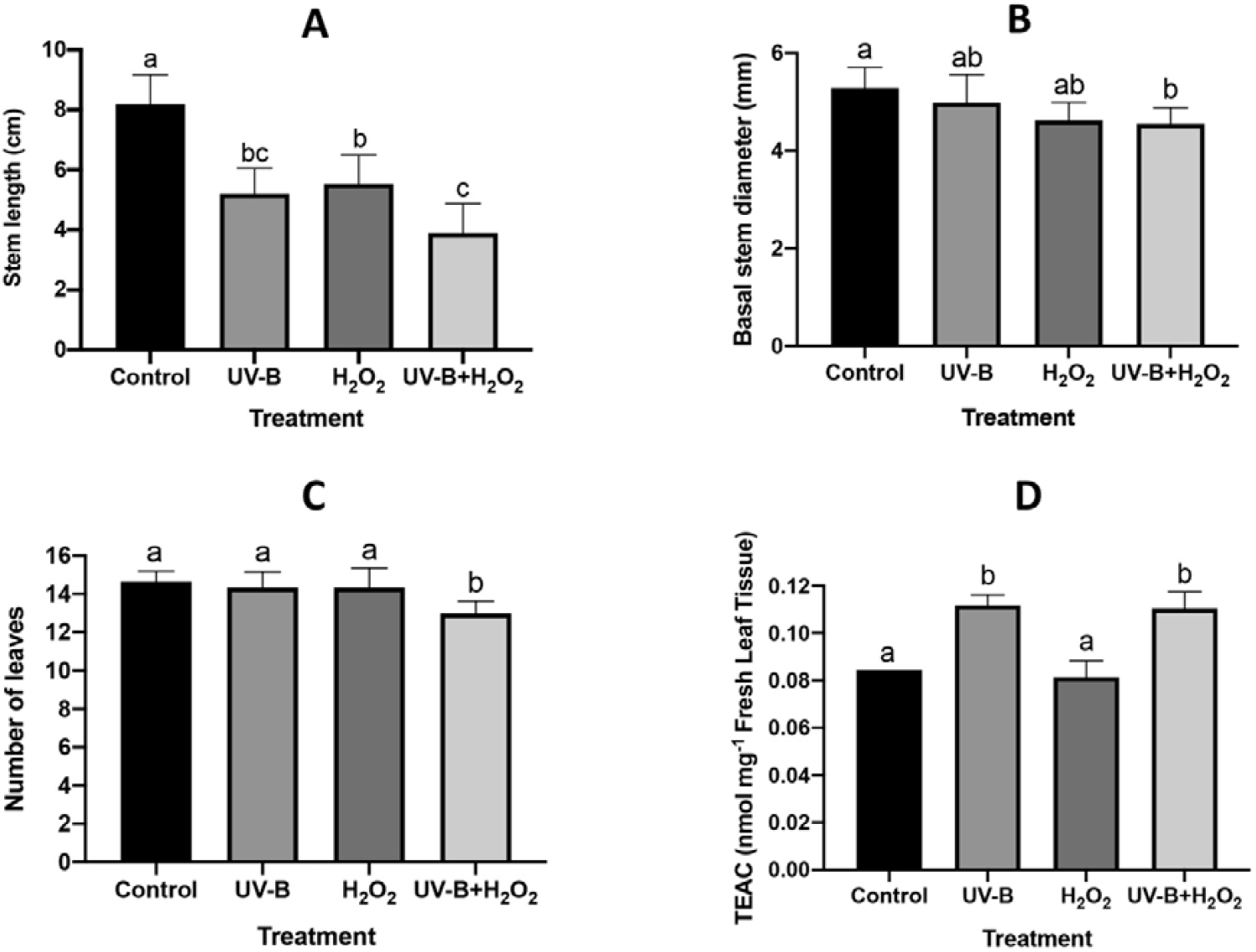
A) Effects of UV-B, H_2_O_2_ and UV-B+H_2_O_2_ on the A) length and B) basal diameter of the stem, C) number of leaves and D) the antioxidant capacity in Trolox equivalents (TEAC) in *N. tabacum* plants after 14 days of pre-treatment. The number of samples per point was n = 6 for morphological data and n = 3 for TEAC data. Different letters indicate significant differences between the treatments using Tukey’s test (P ≤ 0.05). The standard deviation of the measurements is shown.

**Figure 2.**
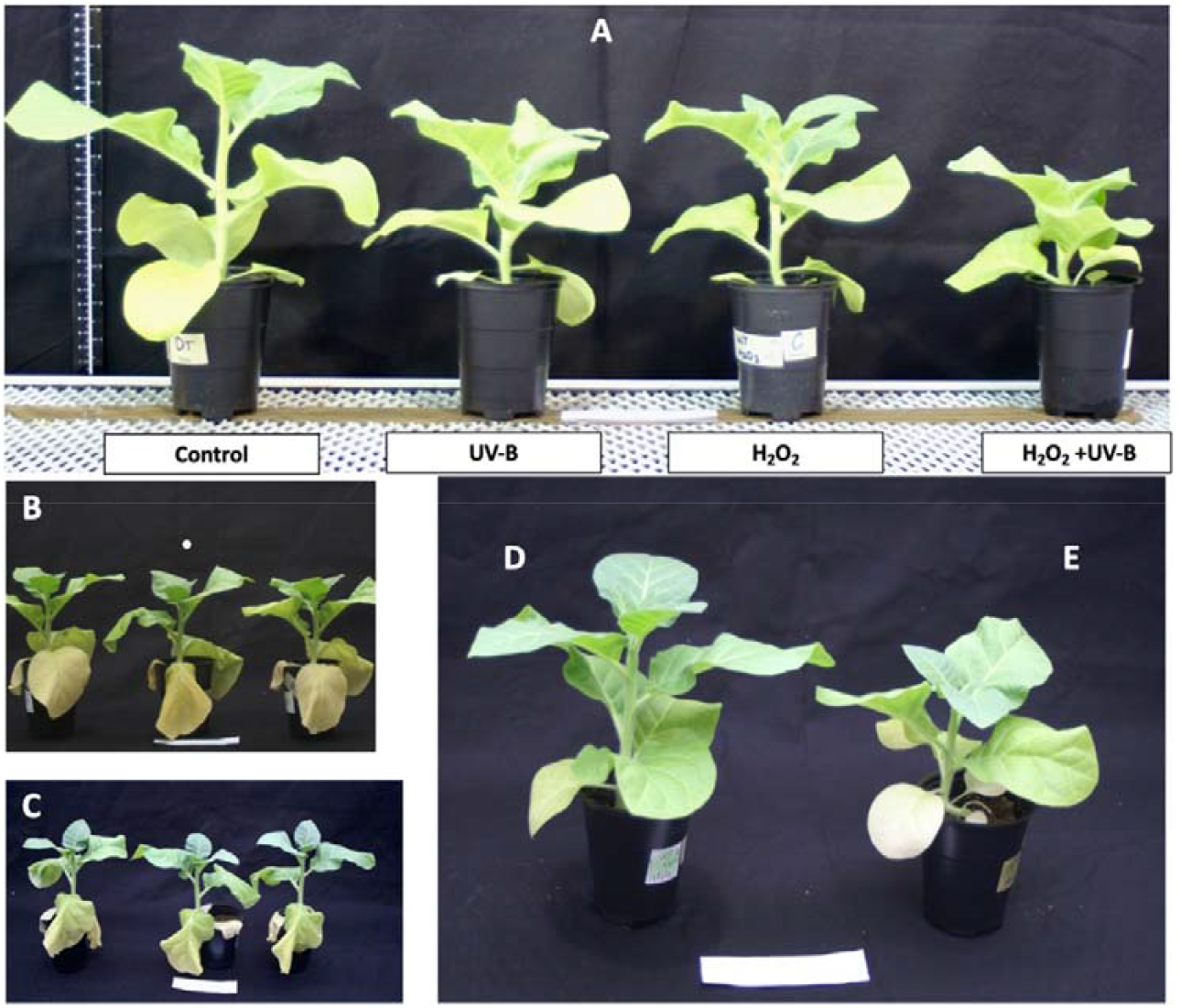
Representative images of A) the pre-treatments after 14 days and wilting after 7 days of drought treatment (days 15-21 of the experiment) in *N. tabacum* plants previously subjected to B) no-pre-treatment, C) UV-B-pre-treatment, D) H_2_O_2_-pre-treatment and E) UV-B+H_2_O_2_-pre-treatment.

The total antioxidant capacity increased in plants subjected to UV-B by 33% compared to the control, whereas no change was observed in H_2_O_2_-treated plants (Fig. 1D). The UV-B+H_2_O_2_ treatment led to increased total antioxidant capacity by 31% compared to the control (Fig. 1D).

The foliar adaxial epidermal flavonol content followed a similar trend, where the UV-B treatment as a single factor increased this parameter compared to the control from day 7 of the experiment and until the end of the pre-treatment exposure. As with the total antioxidative capacity, the H_2_O_2_ treatment did not lead to any changes in the flavonol content throughout the 14 days of exposure (Table 1). For the combined treatment, the difference was significant from hour 28 after the start of the exposure and was maintained throughout the 14 days (Table 1). The combined treatment induced significantly higher leaf epidermal flavonol content between days 5 and 12 when compared to the UV-B treatment.

**Table 1.**
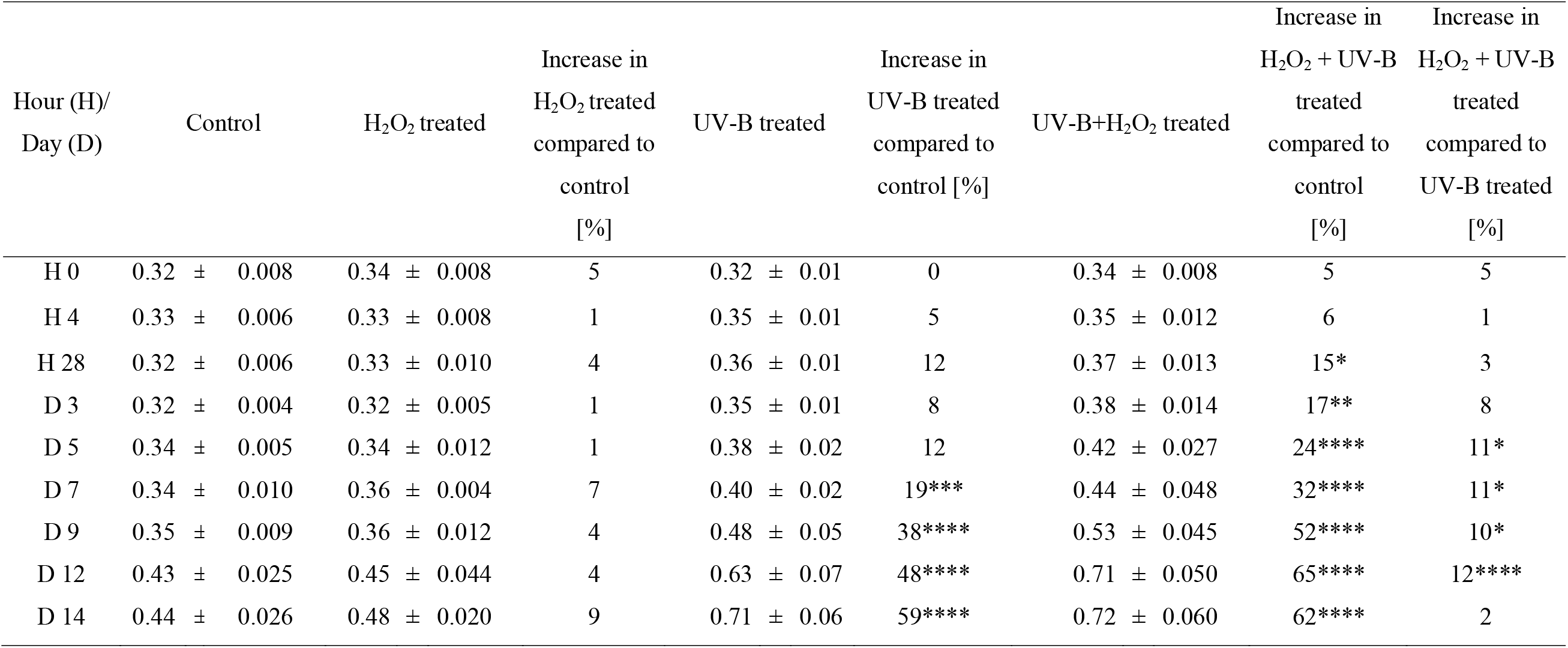
Leaf adaxial epidermal flavonoid content at 0, 4 and 28 hours and at days 3, 5, 7, 9, 12 and 14 in *N. tabacum* leaves cultivated under pre-treatment with UV-B, H_2_O_2,_ or UV-B + H_2_O_2_, or in the absence of pre-treatment. Data are means of n=6 measurements ± standard deviation. Significant differences were determined among the means using Tukey’s test at p < 0.05, 0.01, 0.001 and 0.0001 which are indicated with *, **, ***, ****, respectively.

| Hour (H)/<br>Day (D) | Control | H<sub>2</sub>O<sub>2</sub> treated | Increase in<br>H<sub>2</sub>O<sub>2</sub> treated<br>compared to<br>control<br>[%] | UV-B treated | Increase in<br>UV-B treated<br>compared to<br>control [%] | UV-B+H<sub>2</sub>O<sub>2</sub> treated | Increase in<br>H<sub>2</sub>O<sub>2</sub> + UV-B<br>treated<br>compared to<br>control<br>[%] | Increase in<br>H<sub>2</sub>O<sub>2</sub> + UV-B<br>treated<br>compared to<br>UV-B treated<br>[%] |
| --- | --- | --- | --- | --- | --- | --- | --- | --- |
| H 0 | 0.32 ± 0.008 | 0.34 ± 0.008 | 5 | 0.32 ± 0.01 | 0 | 0.34 ± 0.008 | 5 | 5 |
| H 4 | 0.33 ± 0.006 | 0.33 ± 0.008 | 1 | 0.35 ± 0.01 | 5 | 0.35 ± 0.012 | 6 | 1 |
| H 28 | 0.32 ± 0.006 | 0.33 ± 0.010 | 4 | 0.36 ± 0.01 | 12 | 0.37 ± 0.013 | 15* | 3 |
| D 3 | 0.32 ± 0.004 | 0.32 ± 0.005 | 1 | 0.35 ± 0.01 | 8 | 0.38 ± 0.014 | 17** | 8 |
| D 5 | 0.34 ± 0.005 | 0.34 ± 0.012 | 1 | 0.38 ± 0.02 | 12 | 0.42 ± 0.027 | 24**** | 11* |
| D 7 | 0.34 ± 0.010 | 0.36 ± 0.004 | 7 | 0.40 ± 0.02 | 19*** | 0.44 ± 0.048 | 32**** | 11* |
| D 9 | 0.35 ± 0.009 | 0.36 ± 0.012 | 4 | 0.48 ± 0.05 | 38**** | 0.53 ± 0.045 | 52**** | 10* |
| D 12 | 0.43 ± 0.025 | 0.45 ± 0.044 | 4 | 0.63 ± 0.07 | 48**** | 0.71 ± 0.050 | 65**** | 12**** |
| D 14 | 0.44 ± 0.026 | 0.48 ± 0.020 | 9 | 0.71 ± 0.06 | 59**** | 0.72 ± 0.060 | 62**** | 2 |

On the last day of exposure, the foliar epidermal content of flavonols was similar between these treatments (Table 1).

### Leaf relative water content and root proline content induced by drought treatment

The first signs of drought stress were visible on day 6 in control plants, which exhibited wilting of their older leaves (as shown for plants on day 7 in Fig. 2B). On day 7, plants exposed to UV-B also already showed signs of wilting (Fig. 2C), whereas plants treated with H_2_O_2_ and UV-B+H_2_O_2_ did not exhibit any visible signs of stress during the experiment (Figs. 2D and 2E). On day 7 of drought treatment (i.e. day 21 of the experiment), the RWC decreased significantly in the control plants by 21.9% (from 95.2% RWC in non-drought-treated plants to 74.3% RWC in drought-treated plants). Correspondingly, the RWC decreased significantly in UV-B-pre-treated plants by 25.5% (from 96.0% RWC in non-drought-treated plants to 71.5% RWC in drought-treated plants) (Table 2). In plants pre-treated with H_2_O_2_ only, the RWC decreased significantly by 6.8% (from 93.3% RWC in non-drought-treated plants to 87.0% RWC in drought-treated plants). The combined UV-B+H_2_O_2_-pre-treatment led to a significant decrease in RWC from 93.5 to 86.2% (i.e. by 7.8%). The proline content in roots did not show any significant changes between treatments in this experiment (Table 2).

**Table 2.** Leaf relative water content (RWC) and root proline content at the end of drought treatment (day 21). Different letters indicate significant differences between the treatments using Tukey’s test (P ≤ 0.05). Data represent means of n=3 measurements ± standard deviation.

| Treatment | Leaf RWC (%) | | Root proline content [$\mu\text{M}/\text{mg}$] | |
| --- | --- | --- | --- | --- |
|  | Control | Drought | Control | Drought |
| Control | $95.17 \pm 1.33^a$ | $74.33 \pm 1.86^b$ | $2.67 \pm 0.74^a$ | $2.94 \pm 0.18^a$ |
| UV-B | $96 \pm 0.63^a$ | $71.50 \pm 1.76^b$ | $2.55 \pm 0.13^a$ | $3.26 \pm 0.54^a$ |
| H<sub>2</sub>O<sub>2</sub> | $93.33 \pm 2.73^a$ | $87 \pm 1.26^c$ | $3.06 \pm 0.98^a$ | $2.87 \pm 0.07^a$ |
| UV-B + H<sub>2</sub>O<sub>2</sub> | $93.50 \pm 2.59^a$ | $86.17 \pm 0.75^c$ | $2.83 \pm 0.40^a$ | $2.94 \pm 0.05^a$ |

### CAT, PAL, POD and SOD activities induced by the drought treatment

PAL activity significantly increased under non-drought conditions in plants of all three pre-treatments compared with pre-treatment controls. Whereas the increase was similar in the UV-B-pre-treated and UV-B + H_2_O_2_-pre-treated plants, H_2_O_2_-pre-treatment led to a significantly higher PAL activity still (Fig. 3A). Drought did not further affect PAL activity in the H_2_O_2_-pre-treatment plants, whereas PAL activity in plants pre-treated with either UV-B or UV-B + H_2_O_2_ was significantly higher under drought conditions. The UV-B-pre-treated plants showed the highest PAL activity during drought conditions, tightly followed by the plants that had received the combined pre-treatment. Interestingly, no pre-treated control plants had a higher PAL activity after drought than the H_2_O_2_-pre-treated plants (Fig. 3A).

**Figure 3.**
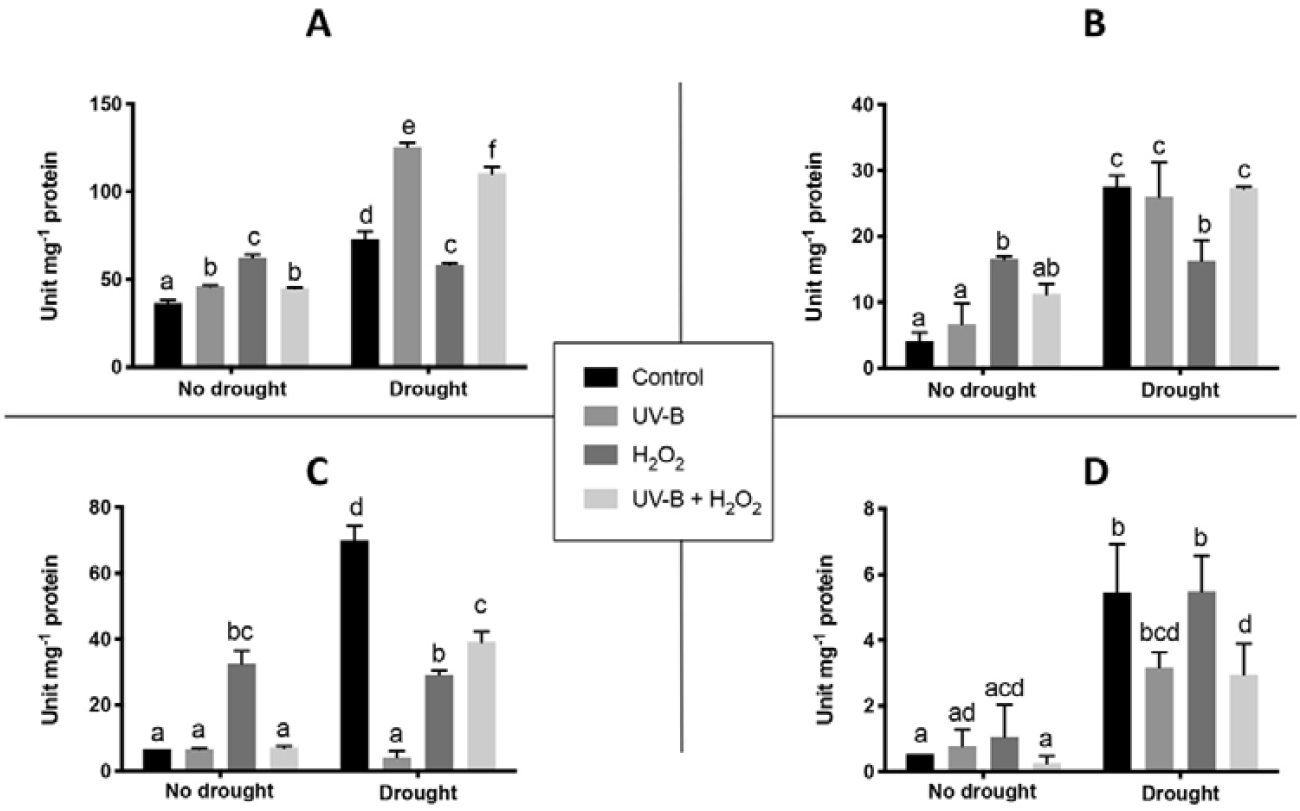
Activities in *N. tabacum* leaves after 21 days of drought treatment of important defense enzymes. A) Phenylalanine ammonia lyase (PAL); B) catalase (CAT); C) Total peroxidase (POD) and D) Superoxide dismutase (SOD). Different letters indicate significant difference between treatments using Tukey’s test (P≤0.05). The standard deviation of the measurements is shown.

The H_2_O_2_-pre-treatment induced significantly higher CAT activity under non-drought conditions than in control plants or UV-B-pre-treated plants (Fig. 3B). Drought again did not alter CAT activity in H_2_O_2_-pre-treated plants, whereas all three other pre-treatments did. Control, UV-B-pre-treated and plants pre-treated with both UV-B and H_2_O_2_ all had a similar CAT activity after drought exposure and was significantly higher than that of the H_2_O_2_-pre-treated plants (Fig. 3B).

Similarly to CAT, H_2_O_2_-pre-treated plants had significantly higher POD activity under non-drought conditions than plants that had experienced any of the other pre-treatments (Fig. 3C). Yet again, drought treatment did not alter POD activity in H_2_O_2_-pre-treated plants. The low POD activity under non-drought conditions in UV-B-treated plants did not change as the result of drought. However, plants exposed to the double pre-treatment exhibited an approximate 4-fold increase in POD activity after drought. Even more pronounced was the increase in control plants of the POD activity when drought was applied, induction being two-fold larger still (Fig. 3C).

Finally, the SOD activity of non-drought exposed plants was low independently of what pre-treatment the plants had been given. Any differences between the pre-treatments were too small to be statistically different (Fig. 3D). In this case, drought did lead to increased SOD activities in plants after all four pre-treatments, the increase being considerably larger in control plants and in H_2_O_2_-pre-treated plants.

### Total antioxidant capacity and flavonol content induced by the drought treatment

Total antioxidant capacity was similar in all pre-treatment groups under non-drought conditions (Fig. 4A). On day 21, drought led to significant increase in the control group and in the UV-B-pre-treated plants. On the other hand, the H_2_O_2_ and UV-B+H_2_O_2_-pre-treated plants did not show any increase between drought and non-drought conditions (Fig. 4A).

**Figure 4.**
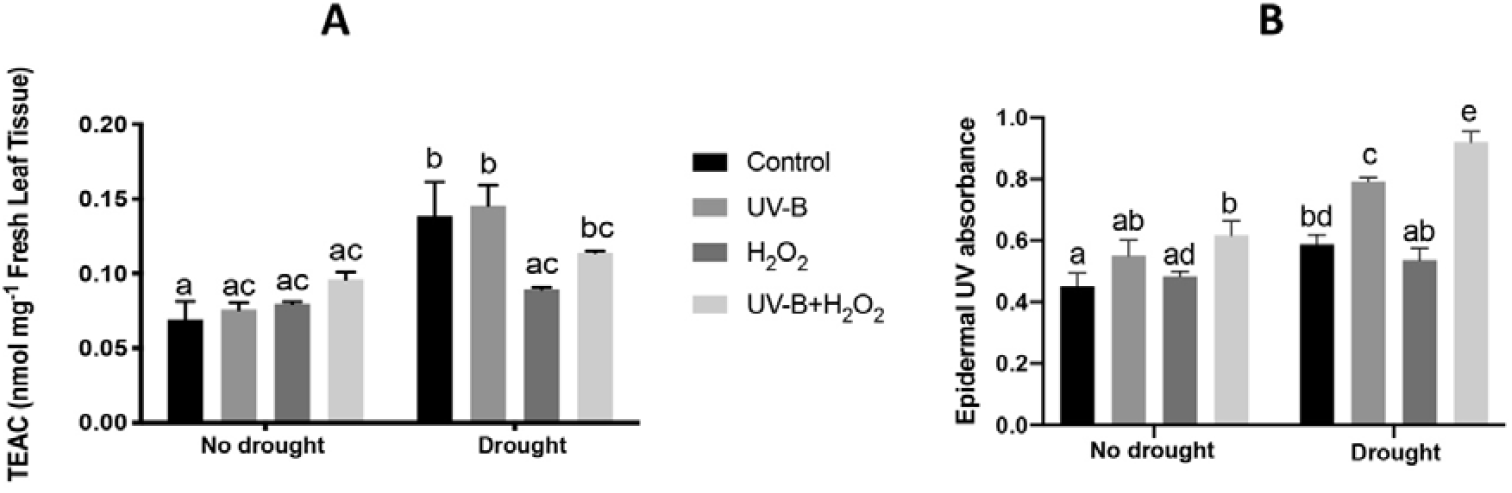
Effects of UV-B-, H_2_O_2_- and UV-B+H_2_O_2_-pre-treatments on *N. tabacum* responses to 7 days of drought treatment (day 21 of the experiment). A) Trolox equivalent antioxidant capacity (TEAC) and B) leaf epidermal flavonol content estimated with Dualex. Different letters indicate significant difference between treatments using Tukey’s test (P≤0.05). The standard deviation of the measurements is shown.

UV-B+H_2_O_2_-pre-treated plants exhibited significantly higher content of leaf epidermal flavonols under non-drought conditions than H_2_O_2_-pre-treated and control plants on day 21 (Fig. 4B). Drought treatment increased the content even more in UV-B+H_2_O_2_-pre-treated plants, which showed the highest levels, followed by the UV-B-treated plants and lastly the non-pre-treated plants (Fig. 4B). Drought treatment did not increase the content of leaf epidermal flavonols in H_2_O_2_-pre-treated plants (Fig. 4B).

### Effects of UV-B, H_2_O_2,_ and drought on transcript accumulation of UVR8, NAC13, BES1 and CHS

Increased transcript accumulation of *UVR8* was only triggered in UV-B- and UV-B+H_2_O_2_-pre-treated plants on day 14 (Fig. 5A). At this time point, UV-B-pretreatment led to a 27-fold increase in the *UVR8* mRNA levels when compared with the corresponding control. Plants pre-treated with UV-B+H_2_O_2_ showed a 63-fold increase in *UVR8* transcript levels (Fig. 5A). On day 21, the *UVR8* transcript abundance had returned to control levels, independently of pre-treatment and whether the plants had been exposed to drought or not.

**Figure 5.**
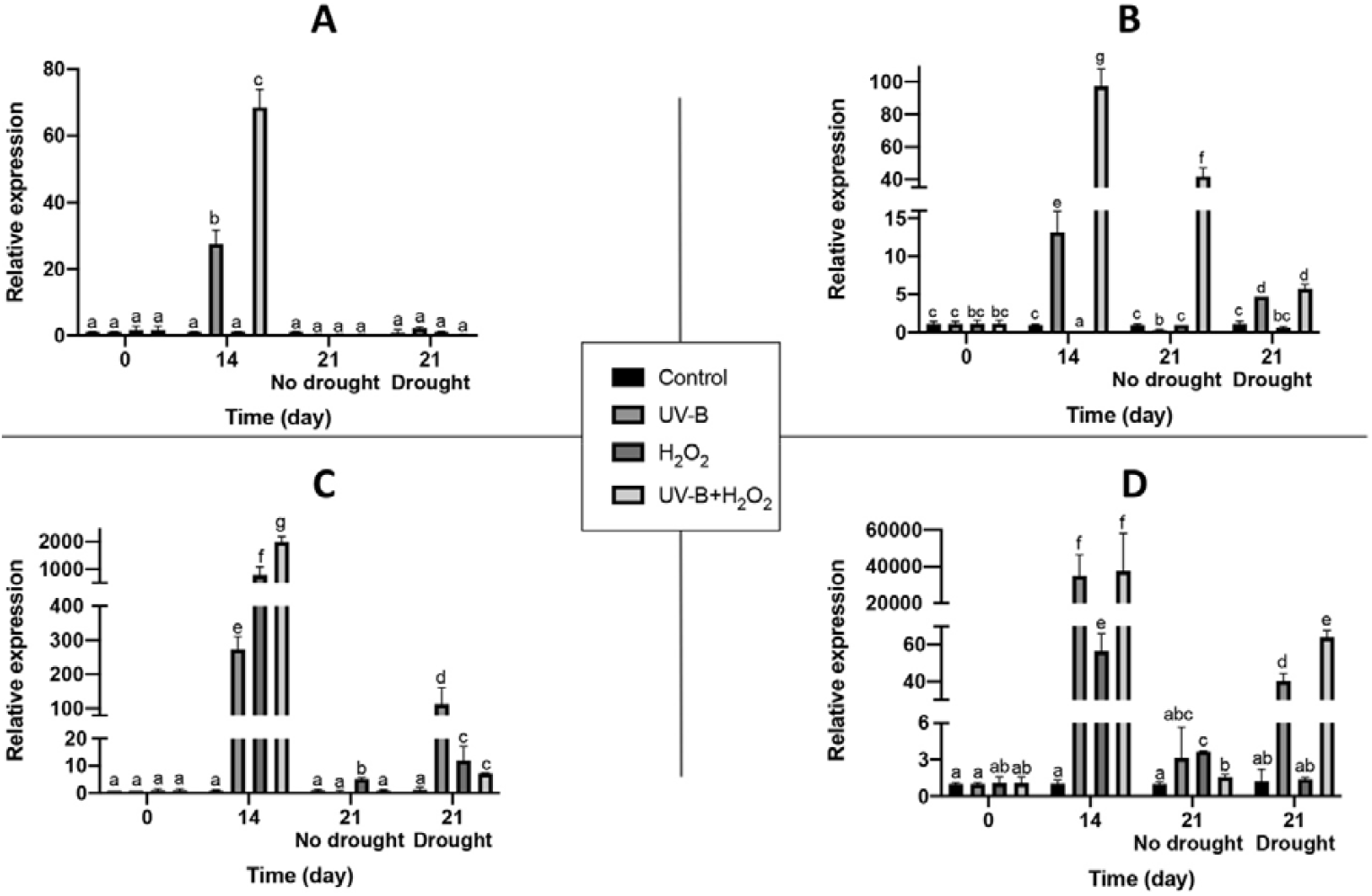
Transcript accumulation for A) *UV RESISTANCE LOCUS 8* (*UVR8*); B) *NAC DOMAIN CONTAINING PROTEIN 13* (*NAC13*); C) *BRI1-EMS-SUPPRESSOR 1* (*BES1*); D) *CHALCONE SYNTHASE* (*CHS*) genes on days 0, 14 and 21 after commencement of the experiment. Pre-treatment was performed for the first 14 days and drought treatment was performed on days 15-21. Relative expression levels were measured in *N. tabacum* leaves using qPCR. Each reaction was performed at least in duplicate in two independent experiments. Error bars represent standard errors. Different letters indicate a significant difference between the treatments using the Tukey’s test (P≤0.05).

For *NAC13*, on day 14 the expression pattern was very similar to that for *UVR8*. UV-B and UV-B+H_2_O_2_ pre-treatments led to transcript accumulation by 13- and 97-fold, respectively (Fig. 5B). In contrast to *UVR8, NAC13* transcript levels were still enhanced (by 41-fold) on day 21 in well-watered UV-B+H_2_O_2_-pre-treated plants. In drought-treated plants on day 21, both UV-B- and UV-B+H_2_O_2_-pre-treated plants had a similar *NAC13* expression about 5-fold above the control level (Fig. 5B).

*BES1* transcript levels increased on day 14 by 267-, 772- and 1956-fold, in plants that had received UV-B-, H_2_O_2_- and UV-B+H_2_O_2_–pre-treatments, respectively (Fig. 5C). On day 21, i.e. after 7 days of drought, *BES1* levels were higher in plants pre-treated with UV-B (100-fold), H_2_O_2_ (12-fold) and UV-B+H_2_O_2_ (7-fold) and exposed to drought than the no-pretreatment control plants (Fig. 5C). On day 21 under no-drought conditions, only the H_2_O_2_-pre-treatment led to increased *BES1* transcript levels compared to the corresponding control (by 5-fold) (Fig. 5C).

On day 14, tobacco plants displayed a clear and significant *CHS* mRNA induction by UV-B-, H_2_O_2_-, and UV-B+H_2_O_2_–pre-treatments by approximately 34000-, 55- and 37000-fold, respectively, compared to the no-pre-treatment control plants. A similar trend was maintained on day 21 after 7 days of subsequent drought treatment for the plants that had been pre-exposed to UV-B-(40-fold) and UV-B+H_2_O_2_-(65-fold) pre-treatments (Fig. 5D). In contrast, on day 21 in no-drought treated plants, the *CHS* transcript levels had almost returned to the basal level, only displaying a 2-4-fold increase compared to the corresponding control in all cases (Fig. 5D).

## Discussion

The increase of plants’ tolerance to stress conditions through the application of physical and chemical factors is accompanied by morphological, biochemical and molecular changes, which are important to study since their interactions determine the desired response. In this study, the effects of UV-B radiation and exogenously applied H_2_O_2_ on a subsequent drought treatment in tobacco plants were studied independently and combined through the assessment of several morphological and physiological parameters.

With regards to the morphological parameters, the shorter stem length exhibited by the UV-B-pre-treated plants (Fig. 1A) agrees with previous studies reporting significant decreases in stem length in different plant species under UV light regimens (Robson, Klem, Urban & Jansen, 2015b; Rodríguez-Calzada et al., 2019; Qian et al., 2020). In contrast, with regards to H_2_O_2_-pre-treated plants, our results differ from other reports that have shown increased biomass and plant height in response to exogenously applied H_2_O_2_ (Ashfaque, 2014; Sun *et al*. 2016; Basal & Szabó 2020). In these previous papers, and in most studies where exogenous H_2_O_2_ have been applied, foliar spraying has been used as the sole application method. In contrast, in our H_2_O_2_-pre-treatment we also apply hydrogen peroxide to roots through irrigation, a difference that might influence the results on plant morphology.

The observed increase in the total antioxidant capacity of UV-B- and UV-B+H_2_O_2_-treated plants at the end of the pre-treatment period (Fig. 1D) could be explained by an increase in non-enzymatic antioxidants, since the foliar epidermal flavonol content was significantly higher in these groups of plants. Furthermore, our data indicate that regarding these two biochemical parameters, plants treated with UV-B+H_2_O_2_ were clearly influenced by the effect of the UV-B treatment rather than by H_2_O_2_. This novel finding highlights the need to study the dominant convergence of UV-B and ROS signaling pathways against other possible pathways triggered by additional factors, for conferring defense responses in plants. The UV-B-induced foliar flavonol content in this experiment (Table 1) is consistent with the fact that a primary mechanism by which plants acclimate to UV exposure is the accumulation of UV-absorbing compounds, such as flavonoids and anthocyanins in leaf epidermal tissue, acting as UV shielding components, but which also are efficient ROS scavengers (Agati & Tattini 2010; Agati *et al*. 2013; Neugart & Schreiner 2018).

The leaf RWC and proline content were measured on day 21 to evaluate differences between those pre-treatments that led to drought tolerance (H_2_O_2_ and UV-B+H_2_O_2_) and those that induced foliar wilting caused by the drought treatment (Control and UV-B). The wilting observations (Fig. 2) were corroborated by RWC measurements (Table 2), the values of which were lower in plants that presented this trait. H_2_O_2_ and UV-B+H_2_O_2_ treatments significantly reduced losses in leaf water content compared to control and UV-B treatments, but the fact that there was no change in proline content suggests that drought stress tolerance was not due to osmotic adjustment related to this osmolyte (Table 2). Similar results in which RWC was increased due to the effect of exogenous H_2_O_2_ have been reported under different drought stress regimens (Ishibashi *et al*. 2011; Sun *et al*. 2016; Basal & Szabó 2020). Thus, our data together with previous studies indicate that a strong defense response of H_2_O_2_ exposure under drought is the maintenance of high levels of leaf RWC, an effect that was maintained in the combined pre-treatment, and which probably has a key role in the tolerance shown by the plants.

CAT is considered a major H_2_O_2_-scavenging enzyme which is mainly active at relatively high H_2_O_2_ concentrations (Anjum *et al*. 2016). Our UV-B treatment did not affect CAT activity in non-drought-treated plants (Fig. 3B), which indicates that the low UV-B doses used did not induce significant changes in the production of H_2_O_2_. Since POD and SOD activities did not increase in the UV-B-pre-treated group either (Figs. 3C and 3D), this also confirms that the treatment did not induce oxidative stress. Increases in CAT, SOD and POD activities by UV-B has previously been reported in experiments using higher daily doses. Priming with 6 kJ m^-2^ of UV-B radiation in rice seedlings induced CAT, as well as SOD and ascorbate peroxidase (APX) activities (Thomas & Puthur 2019). In another study, exposure of *N. tabacum* plants to high (13.6 kJ m^-2^ d^-1^) biologically effective UV-B radiation led to increases in peroxidase defense, specifically APX, and of the chloroplast-localized Fe-SOD (Majer, Czégény, Sándor, Dix & Hideg, 2014). Rácz, Hideg & Czégény (2018) also reported that leaf acclimation to UV-B, corresponding to a 7.7 kJ m^−2^ d^−1^ biologically effective dose, is carried out via a selective activation of POD isoforms. Our data showed that UV-B exposure specifically increased PAL activity (Fig. 3A) and stimulated the synthesis of foliar epidermal flavonols (Table 1) in the absence of oxidative stress.

The high level of CAT activity in H_2_O_2_-pre-treated tobacco plants under non-drought conditions (Fig. 3B) indicates that the application of this elicitor either led to uptake of this uncharged molecule by the plant, or to induction of ROS production. Moreover, the CAT activity remained at the same level in drought-treated plants, which shows that a durable post-application effect was induced by the exogenous H_2_O_2._ The PAL and POD activities of H_2_O_2_-pre-treated tobacco plants showed the same behavior (Figs. 3A and 3C). These results suggest that activation of the enzymatic antioxidant machinery due to H_2_O_2_ pretreatment induced the drought tolerance that these plants did obtain.

Other studies have also concluded that the capacity for osmotic adjustment via up-regulating antioxidant enzymes influences the drought tolerance due to elicitation by H_2_O_2_. Sun et al. (2016) found that drought tolerance of cucumber seedlings was improved by foliar application of 1.5 mM H_2_O_2_. In their study, the activity of CAT was significantly increased under severe drought conditions but not under moderate drought, contrary to the SOD and POD activities, which were significantly increased by the elicitor under medium drought-stress conditions. Similarly, exogenous soil application of 5 mmol/L H_2_O_2_ improved water stress tolerance of rice seedlings at the same time as CAT activity remained unaltered compared with the control. Ascorbate peroxidase (APX) and guaiacol peroxidase (GPOX) activities were however significantly increased (Sohag et al., 2020).

With regards to UV-B+H_2_O_2_-pretreated plants, the enzymatic activities of PAL, CAT and SOD (Figs. 3A, 3B and 3D) followed a similar trend as those of the UV-B-pretreated plants, again confirming the strong influence of UV-B factor in these plants. Interestingly, with regards to POD (Fig. 3C), neither UV-B-pretreatment nor H_2_O_2_-pretreatment led to any increases in activities as the result of drought treatment. However, the combined UV-B+H_2_O_2_-pretreatment led to a synergistic increase in POD activity requiring both pretreatment factors to be present for induction of this response.

To further study the influence of UV-B under the different treatments, we measured changes in transcript accumulation of a set of UV-B marker genes. The UV-B photoreceptor UVR8 (Jenkins, 2014) mediates responses to UV-B through interactions with specific signaling components and transcription factors which ultimately lead to the biosynthesis of phenolic compounds, such as flavonoids (Hideg, Jansen & Strid, 2013; Hideg & Strid, 2017; Jenkins, 2014; Liang, Yang & Liu, 2019; Rai et al., 2020). Transcript accumulation of *UVR8* was strongly increased in UV-B-treated tobacco plants on day 14. Interestingly, *UVR8* in Arabidopsis does not appear to be regulated neither at the transcript nor at the protein level (Morales *et al*. 2013; Jenkins, 2014). Thus, transcriptional regulation of *UVR8* by UV-B appears to be species-dependent. Our data further indicate that ROS and UV-B signaling interact to regulate *UVR8* expression in tobacco (Fig 5A). A similar synergistic effect on the mRNA abundance was also observed on day 14 for the *NAC13* and *BES1* genes, given the higher transcript accumulation after UV-B+H_2_O_2_ pre-treatment as compared to plants pre-treated with UV-B or H_2_O_2_ alone (Figs 5B and C). Thus, convergence of UV-B and ROS signaling impacts on the expression of genes involved in multiple pathways.

The increase of *NAC13* transcript levels in UV-B- and UV-B+H_2_O_2_-pre-treated plants on day 14 (Fig. 5B) indicates that *NAC13* in tobacco is induced by low UV-B doses. Although the transcript content decreased between days 14 and 21, particularly in UV-B-pre-treated plants, our results suggest that drought helped to maintain an elevated *NAC13* expression in these plants. *NAC13* appears to be an important gene to mediate diverse stress responses in plants. *NAC13* expression is induced by UV-B in *Arabidopsis* independently of the UVR8 photoreceptor (O’Hara et al., 2019) and is also induced by salt stress in poplar (Zhang et al., 2019), and by drought in broomcorn millet (Shan et al., 2020). In our study, however, drought on its own did not induce *NAC13* expression in tobacco (Fig. 5B).

*BES1* is a transcription factor involved in brassinosteroid (BR) signaling that, when activated, regulates expression of thousands of BR-responsive genes (Clouse, 2011). In response to UV-B, the expression of *BES1* in *Arabidopsis* appears to be dependent on the nature of the UV-B treatment. While UV-B levels in natural sunlight (1.5 μmol m^-2^ s^-1^ for 6 hours) did not affect *BES1* transcript accumulation in wild type plants (Rai et al., 2020), exposure to UV-B stress provided by broad-band UV (5 μmol m^-2^ s^-1^ for 8 hours for 10 days) lowers *BES1* transcript levels (Liang et al., 2020). Here we show that low level UV-B-pre-treatment led to increased *BES1* transcript levels in tobacco plants (Fig. 5C). Furthermore, plants exposed to H_2_O_2_- or combined UV-B+H_2_O_2_-pre-treatments showed an even greater increase in *BES1* mRNA, revealing an important role for H_2_O_2_ in induction of *BES1* expression. These data correlate with recent studies that have shown a positive H_2_O_2_ regulation of BR signaling. Tian et al. (2018) uncovered a critical role of H_2_O_2_ in BR signaling in *Arabidopsis* plants through the oxidation of the BES1 homologous transcription factor BRASSINAZOLE-RESISTANT1 (BZR1), leading to its interaction with key regulators in cell growth and the auxin-signaling pathway. In our study, *BES1* transcript abundance decreased from day 14 to 21 in well-watered plants exposed to all three pre-treatments. However, significant amounts of *BES1* mRNA were maintained on day 21 under drought conditions in plants subjected to the three different pre-treatments. These findings indicate that *BES1* in tobacco is transcriptionally regulated by low UV-B doses and exogenous H_2_O_2_. Drought obviously also influences *BES1* expression induced by UV-B and H_2_O_2_ pre-treatment, although drought itself could not regulate BES1 transcription.

Flavonoids have long been known to have functional roles in absorbing UV light and to function as antioxidants (Agati & Tattini, 2013; Hideg & Strid, 2017). Furthermore, many research groups have reported that flavonoids have a function in defense against drought, when acting as free radical scavengers (Ramakrishna & Ravishankar, 2011; Nakabayashi et al., 2014; Kumar et al., 2020). Our data is in agreement with previous research which showed that UV-B enhances accumulation of *CHS* transcripts, representing first committed enzyme of the flavonoid biosynthesis pathway (Brown & Jenkins, 2008; Rodríguez-Calzada et al., 2019; Rai et al,. 2020; Santin et al., 2020). It is well known that H_2_O_2_ plays a role in regulation of genes associated with phenolic compund biosynthesis, of which CHS is one (Nyathi & Baker, 2006). Our data also show that drought amplified the expression pattern induced particularly by UV-B-pre-treatment in tobacco, a result that coincides with increased levels of epidermal flavonoids (Fig. 4B). Thus, our data strengthens the notion that flavonoids have an important role in inducing drought tolerance and mitigating oxidative stress. Our data also demonstrate a pre-treatment-dependent response to drought for *NAC13, BES1* and *CHS* transcript accumulation.

Thus, in Fig. 6 we summarize the effects of the three environmental cues UV-B light, oxidative pressure (H_2_O_2_-pre-treatment) and drought and their mechanistic interplay. The UV-B-pre-treatment increased the expression of all four molecular markers in tobacco, whereas drought alone did not alter gene expression (Figs. 5 & 6). H_2_O_2_-pre-treatment does not lead to expression of *UVR8* or *NAC13* (Figs. 5A, 5B & 6), but induces transcription of *BES1* and *CHS* (Fig. 5C, 5D & 6). Furthermore, H_2_O_2_ potentiates the UV-B-induced expression of *BES1* and *CHS* (Fig. 5C, 5D & 6). Drought also prolongs the effect of UV-B-pre-treatment on *NAC13, BES1*, and *CHS* expression (Fig. 5B, 5C, 5D & 6).

**Figure 6.**
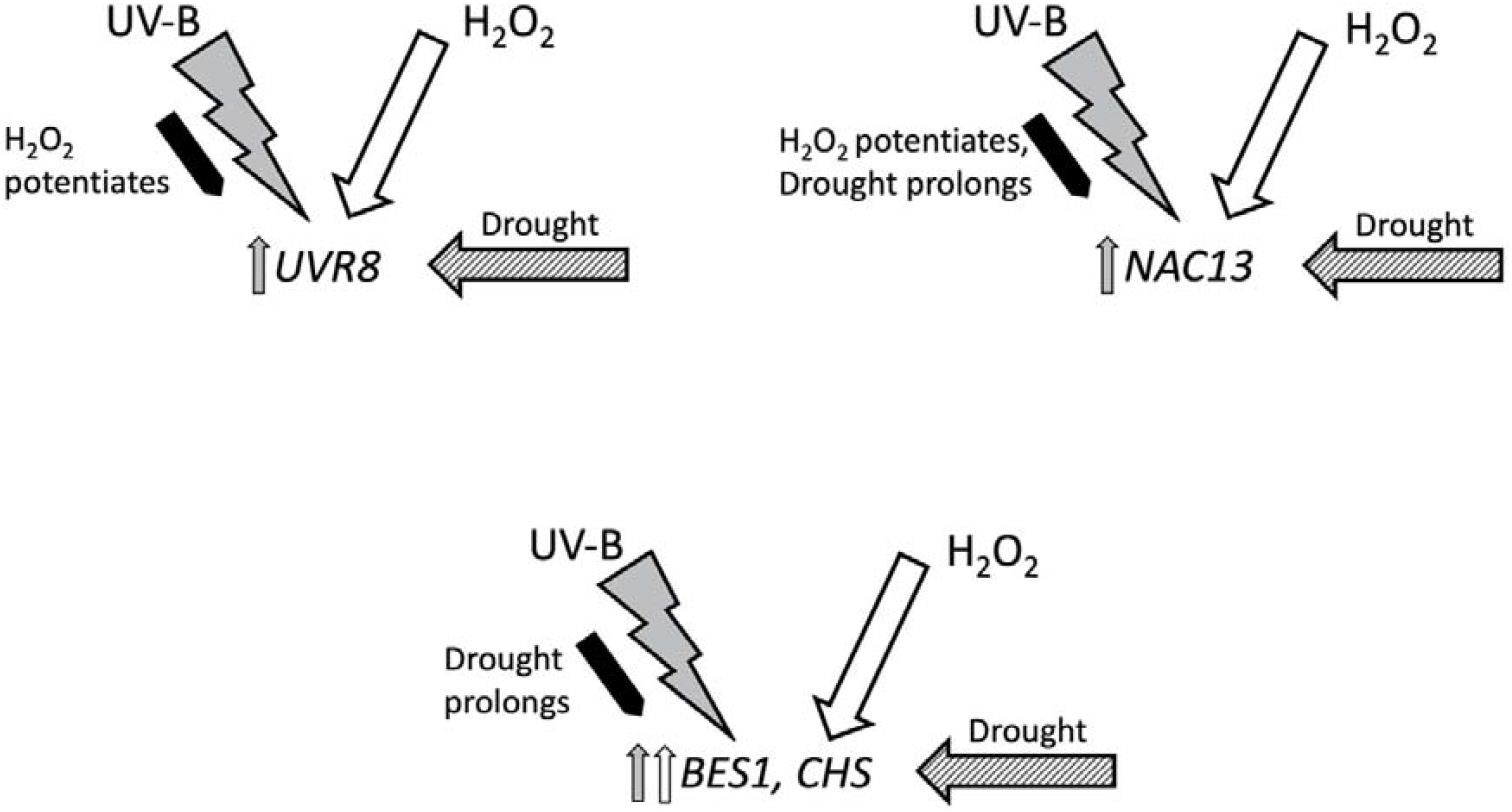
Schematic representation of the effects of UV-B- and H_2_O_2_-pre-treatments and subsequent drought treatment on the expression of *UVR8, NAC13, BES1*, and *CHS* in tobacco leaves. Upward arrows indicate induced expression by a given treatment. Black filled arrows show interactive effects between UV-B and H_2_O_2_-pre-treatments and UV-B-pre-treatment and drought conditions.

To conclude, this study highlights the potential of using a combination of factors to provide resistance to drought which could become a useful tool to for instance reduce water use in agriculture in a current challenging environmental scenario. A key aspect in elicitation with controlled elicitation will be further studies of the biochemical and molecular plant responses toward these factors in order to find specific conditions under which the plants show the desired responses.

## Acknowledgements

This research was supported by research grants to ÅS from the Knowledge Foundation (kks.se; grant #20130164), and the Swedish Research Council Formas (formas.se/en; grant #942-2015-516). The project was also supported by the Faculty for Business, Science and Technology at Örebro University. Moreover, this research was partially supported by SEP-CONACyT (Ciencia basica 2016, grant 283259). D.S-d acknowledges the doctorate scholarship 707895 and travel grant given by Consejo Nacional de Ciencia y Tecnología (CONACyT).

The project was supported by the Faculty for Business, Science and Technology at Örebro University. D.S-d acknowledges the doctorate scholarship 707895 and travel grant given by Consejo Nacional de Ciencia y Tecnología (CONACyT).

## Notes

**Funding**: This research was supported by research grants to ÅS from the Knowledge Foundation (kks.se; grant #20130164), and the Swedish Research Council Formas (formas.se/en; grant #942-2015-516). Moreover, this research was partially supported by SEP-CONACyT (Ciencia basica 2016, grant 283259).

### Competing Interest Statement

The authors have declared no competing interest.

