## Supplementary material for "UV-B exposure and exogenous hydrogen peroxide application leads to cross-tolerance toward drought in *Nicotiana tabacum* L": Suppl.Info

to

**Supplementary Table S1**. Concentrations of macro- and micronutrients in the substrate and watering solutions.

| **NUTRIENT** | **Peat Substrate [ppm]** | **Hoagland Solution, 25% [ppm]** |
|---|---|---|
| Nitrogen (N) | 140 | 52.5 |
| Phosphorous (P) | 70 | 7.75 |
| Potassium (K) | 149 | 58.75 |
| Magnesium (Mg) | 249 | 12 |
| Sulphur (S) | 76 | 16 |
| Calcium (Ca) | 2186 | 50 |
| Iron (Fe) | 0.9 | 0.25 to 1.25 |
| Manganese (Mn) | 1.6 | 0.125 |
| Cupper (Cu) | 1.2 | 0.005 |
| Zinc (Zn) | 0.4 | 0.0125 |
| Barium (B) | 0.3 | 0.125 |
| Molybdenum (Mo) | 2.0 | 0.0025 |
